# Nano-silver and forskolin regulate rice seed germination mediated by *OsPIP2-1*

**DOI:** 10.1101/2023.04.20.537739

**Authors:** Bingxian Chen, Yuanxuan Peng, Qi Zhang, Zhongjian Chen, Hongmei Li, Jun Liu

## Abstract

As timely germination of seeds is crucial to crop yield and quality, the low germination percentage for direct seeding in rice and pre-harvest sprouting during late stage of maturity have extremely adverse effects on rice production. Herein, we found that nano-silver inhibited rice seed germination while forskolin promoted this process. Germination assay showed that 20 μM AgNPs and 0.5 μM forskolin could be the optimal concentrations regulating rice seed germination. At the molecular level, an aquaporin gene, *OsPIP2-1*, was most highly expressed among those homologous genes during rice seed germination, and was decreased by AgNPs and increased by forskolin. Meanwhile, the activity of α-amylase and its genes expression were also inhibited by AgNPs and promoted by forskolin. Next, the germination percentage of *OsPIP2-1* overexpressing seeds was significantly higher than that of wild-type, and that of *OsPIP2-1* knockout seeds was dramatically reduced. Furthermore, compared with the wild type, α-amylase activity and the transcription of *OsAmy3C* and *OsAmy3E* were increased in *OsPIP2-1* overexpressing seeds and decreased in *OsPIP2-1* knockout seeds, indicating that *OsPIP2-1* positively regulates α-amylase. Interestingly, seed germination, α-amylase activity and its genes expression in *OsPIP2-1* overexpressed and knockdout lines could also be inhibited by nano-silver and promoted by forskolin. In conclusion, we propose that nano-silver and forskolin regulated seed germination by modulating the expression of *OsPIP2-1* and thus α-amylase. Our results may bring new application of nano-silver and forskolin in rice cultivation.

## INTRODUCTION

Aquaporin is a water-channel protein that selectively and efficiently transports water molecules across plant intracellular membranes or the plasma membrane (Maurel et al., 2008). Each aquaporin monomer comprises six transmembrane helices and five hydrophilic sequences, among which the B and E sequences contain a repeated and highly conserved tripeptide, namely NPA (Asn-Pro-Ala). The NPA has the hour-glass structure, and contribute to the regulation of aquaporin activity (Törnroth-Horsefield et al.,2006). The two NPA sequences of the B and E rings overlap to form a narrow transmembrane water pore that transports water molecules bidirectionally (Gomes et al.,2009). Aquaporins are encoded by multiple genes and have many subfamilies. Arabidopsis has 35 aquaporin genes (Johanson et al.,2001), and rice has 33 (Sakurai et al.,2005). Bryophytes have the most aquaporin types, followed by dicotyledons and then monocotyledons. Monocotyledons have four types of aquaporins, namely those in the plasma membrane, vacuoles, Nod26-like and small molecular pores (Johanson et al.,2001).

In plants, aquaporins play roles in seed germination, transpiration, photosynthesis, stomatal regulation and stress responses mainly by regulating water transport. For example, 70–80% of water in roots is transported through aquaporins (Zhang and Du, 2005). Certain metal ions can effectively inhibit aquaporin activity and block water transport across a membrane. For example, the binding of Hg^2+^ to conserved cysteine residues proximal to the NAP repeats alters the conformation of the aquaporin, thereby blocking the pore and inhibiting water transport (Esteva-Font et al.,2016). The ion Ag+ inhibits aquaporin even more efficiently, as it binds cysteine as well as histidine residues to alter aquaporin conformation (De Almeida et al., 2017). In animal and human cells, the small-molecule natural product forskolin increases aquaporin activity by regulating cAMP-dependent protein kinase activity, thereby increasing the membrane permeability to water and promoting the absorption of water by cells (Skowronska et al., 2015). Plant aquaporin α-TIP can be phosphorylated in oocytes by animal PKA to modulate the water transport activity. To activate PKA activity, forskolin is essential (Maurel et al., 1997).

Water absorption by seeds is a prerequisite for germination. Water content in seeds changes dramatically during imbibition to drive radicle extension after its emergence from the seed coat to complete germination (Footitt et al., 2019). Absorption of water by germinating seeds occurs in three stages, the first of which is a rapid physical rather than metabolic process that can take place in either living or dead seeds. The second stage is called the stagnant (or plateau) stage, when the matrix in the seed is fully hydrated but vacuoles and new protoplasm have not yet formed. After a period of time, the seeds immediately enter the rapid water absorption period again, during which stored substances are mobilized, transcription and translation are substantially upregulated, and DNA synthesis and cell division increase with a concomitant increase in cellular respiration (Bewley et al., 2012).

Speculation remains concerning the role of aquaporins in the water absorption process during seed germination. It has been established that inhibition of aquaporin function delays the completion of seed germination (Vander Willigen et al., 2006). Although several aquaporins are expressed in seeds (Vander Willigen et al., 2006; Gattolin et al., 2011), it remains unknown whether they help regulate germination. In rice (*Oryza sativa*), aquaporin genes *OsPIP1-1*, *OsPIP1-3* and *OsPIP2-3* are expressed in the embryo and seedlings and participate in water regulation by responding to NO signaling, which affects germination rate (Liu et al., 2007; 2013). During seed germination of *Arabidopsis thaliana*, *AtTIP3-1* and *AtTIP3-2* act antagonistically to modulate the response to signaling via the phytohormone abscisic acid, with *AtTIP3-1* being a positive and *AtTIP3-2* a negative regulator. A third isoform encoded by *AtTIP4-1* is normally expressed upon completion of germination (Footitt et al., 2019).

It is well known that high germination percentage is the key to stable rice yield. On the other hand, precocious germination on the ear of rice before harvest also reduces rice yield and quality. In order to solve the above problems, we selected a large number of compounds and plant extracts as seed germination regulators, and finally chose two substances which could effectively affect the germination of rice seeds. We found that nano-silver could inhibit pre-harvest sprouting, and forskolin could accelerate the germination of rice seeds. The mechanism, however, is still unclear. In this study, the role of aquaporin gene OsPIP2-1 in rice seed germination was confirmed by quantitative reverse transcription-PCR (qRT-PCR), in situ hybridization, western blotting, gene overexpression, gene knockout and physiological methods. The underlying mechanism revealed that nano-silver and forskolin regulate rice seed germination by inhibiting and promoting the expression of *OsPIP2-1* and α-amylase, respectively. The results not only clarified the function of *OsPIP2-1* in rice seed germination, but also provided a theoretical basis for the application of nano-silver and forskolin in rice cultivation.

## RESULTS

### AgNPs negatively regulate germination whereas forskolin positively regulates germination as well as water absorption by rice seeds

Whether nano-silver (AgNPs) affect rice seed germination has not been reported. We first characterized the AgNPs reagent with respect to chemical composition and size distribution. Transmission electron microscopy revealed that the AgNPs were roughly spherical, with an average particle size of 44.58 nm (range, 19.9–66.9 nm; Figure 1a-b). SEM-EDX analyses confirmed the expected chemical composition of the AgNPs sample (Figure 1c), and the spectra revealed the presence of Ag in the sample (Figure 1d, Figure S1).

**Figure 1.**
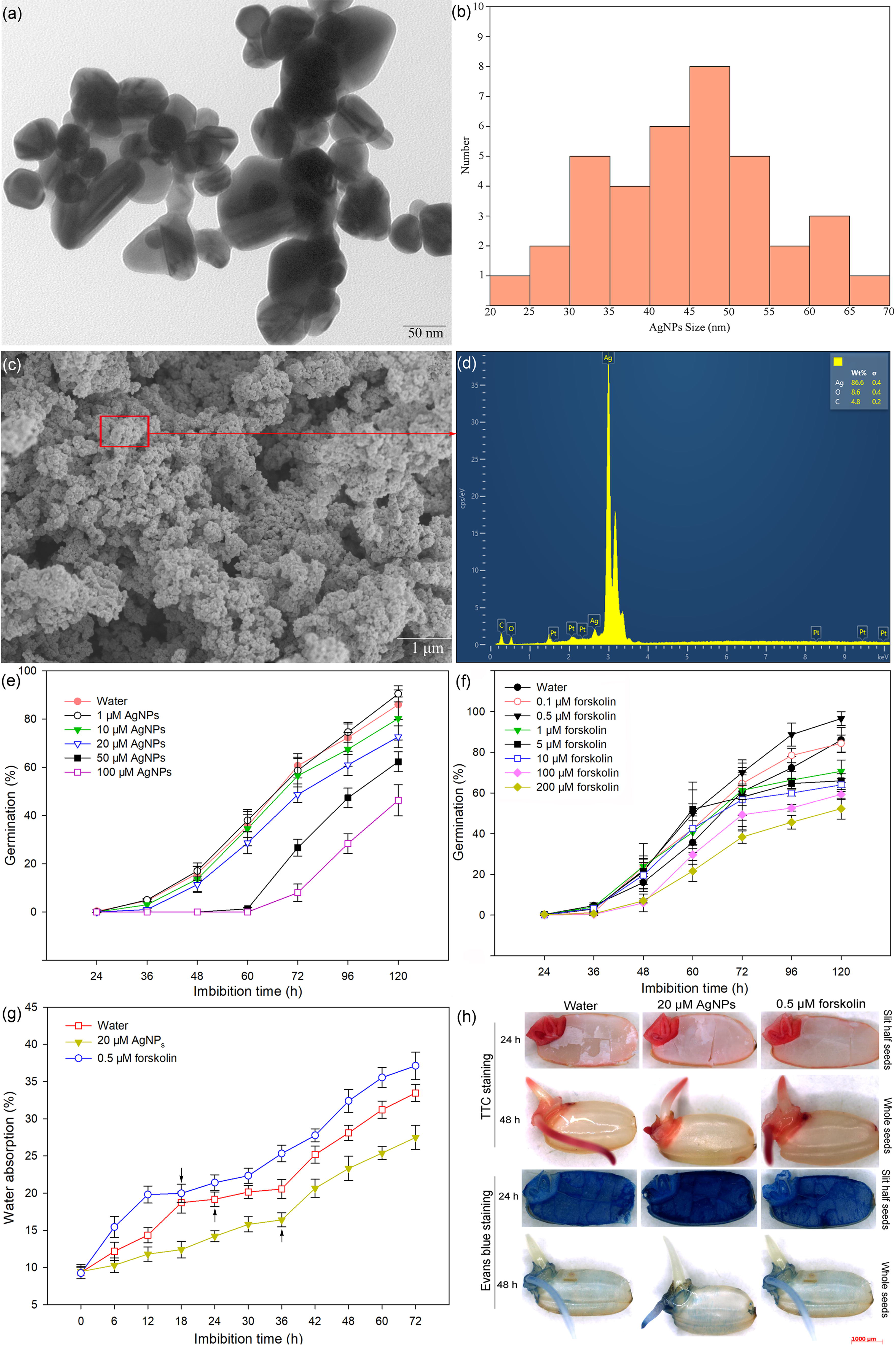
Characterization of nano-silver and rice seed germination. (a) Representative transmission electron micrograph of AgNPs. Scale bar, 30 nm. (b) Size distribution of AgNPs. (c) Representative SEM image of AgNPs. Scale bar, 1 μm. (d) EDX elemental mapping of AgNPs. (e) Germination time curves for rice seeds imbibed in 1, 10, 20, 50, or 100 μM AgNPs or water. (f) Germination time curves for rice seeds imbibed in 0.1, 0.5, 1, 5, 10, 100, or 200 μM forskolin or water. (g) Water absorption curves for rice seeds imbibed in water or 20 μM AgNPs or 0.5 μM forskolin. The black arrow shows the point at which the seed begins to germinate. (h) Effects of AgNPs and forskolin on seed viability.

Rice seeds began to germinate after 24 h of imbibition in water (as the control), and then the germination percentage gradually increased, reaching 36% at 60 h and 86% at 120 h (Figure 1e-f). As shown in Figure 1e, the lowest concentrations of AgNPs tested (1 and 10 µM) did not significantly affect seed germination. However, higher concentrations of AgNPs delayed germination. At 20 μM, AgNPs significantly delayed seed germination but did not harm seedlings; concentrations of 50 and 100 μM AgNPs also significantly delayed germination but resulted in blackening of the germinated radicle and seedling root, presumably owing to the toxicity to roots of Ag at relatively high concentrations (Figure S2). Therefore, 20 μM AgNPs was selected for use in subsequent experiments.

The germination curve for seeds imbibed in 0.1 μM forskolin was similar to that in water. With 0.5 μM forskolin, however, the germination percentage was 50% at 60 h and 97% at 120 h, values that were higher than that of the control (Figure 1f). Indeed, forskolin at 1 μM, 5 μM or 10 μM promoted germination in the early stage of imbibition (before 72 h), but these concentrations inhibited germination after 72 h. Furthermore, forskolin at 100 μM or 200 μM delayed germination (Figure 1f, Figure S2). These results demonstrated that 0.5 μM forskolin could promote rice seed germination and therefore was chosen for use in subsequent experiments.

Water uptake is essential to for seed germination. To investigate the effects of AgNPs and forskolin on water absorption of rice seeds, we monitored changes in water absorption of seeds imbibed in 20 μM AgNPs or 0.5 μM forskolin (Figure 1g). For seeds imbibed in water (control), water absorption increased rapidly up to 18 h, then increased more slowly from 18 h to 36 h to a maximum of 20% by weight, with seed water content remaining at 20%. This stage constitutes the plateau phase of water uptake by seeds. After 36 h, water absorption began to increase rapidly concomitantly with the onset of germination. At 20 μM, AgNPs reduced the time required to reach the plateau period by 6 h, i.e., from 36 h (control) to 30 h. However, 0.5 μM forskolin increased the rate of water absorption at each time point, and reduced the time to reach the onset of the plateau period, i.e., the period began at 12 h vs. 36 h for the control.

TTC (2,3,5-tryphenyl tetrazolium chloride) staining and Evans blue staining were used to judge seed viability. TTC dyes viable tissues red, whereas Evans blue dyes non-viable seeds blue (Pradhan et al., 2022). As shown in Figure 1h, compared with the control, AgNPs and forskolin had almost no effect on embryo color, indicating that any regulation of germination by AgNPs or forskolin did not affect embryo viability.

### The aquaporin genes *OsPIP2-1* and *OsTIP1-1* are expressed at the highest levels, and *OsPIP2-1* is decreased by AgNPs and increased by forskolin

Aquaporins play an important role in water absorption during seed germination, therefore, the expression of aquaporins of rice seeds imbibed in water were analyzed. Rice aquaporin is a multi-gene family, which can be divided into PIP, TIP, NIP and SIP according to their evolutionary relationship (Figure 2a). Next, we assessed the expression of 34 aquaporin genes, among which mRNAs for 5 genes were not detected when rice seeds were imbibed in water. Notably, *OsPIP2-1*, *OsTIP1-1* and *OsPIP1-1* were up-regulated compared with the other aquaporin homologs (Figure 2 b-c). Based on these results, *OsPIP2-1* and *OsTIP1-1*—as the mostly highly expressed aquaporin genes—were selected for analysis in subsequent studies.

**Figure 2.**
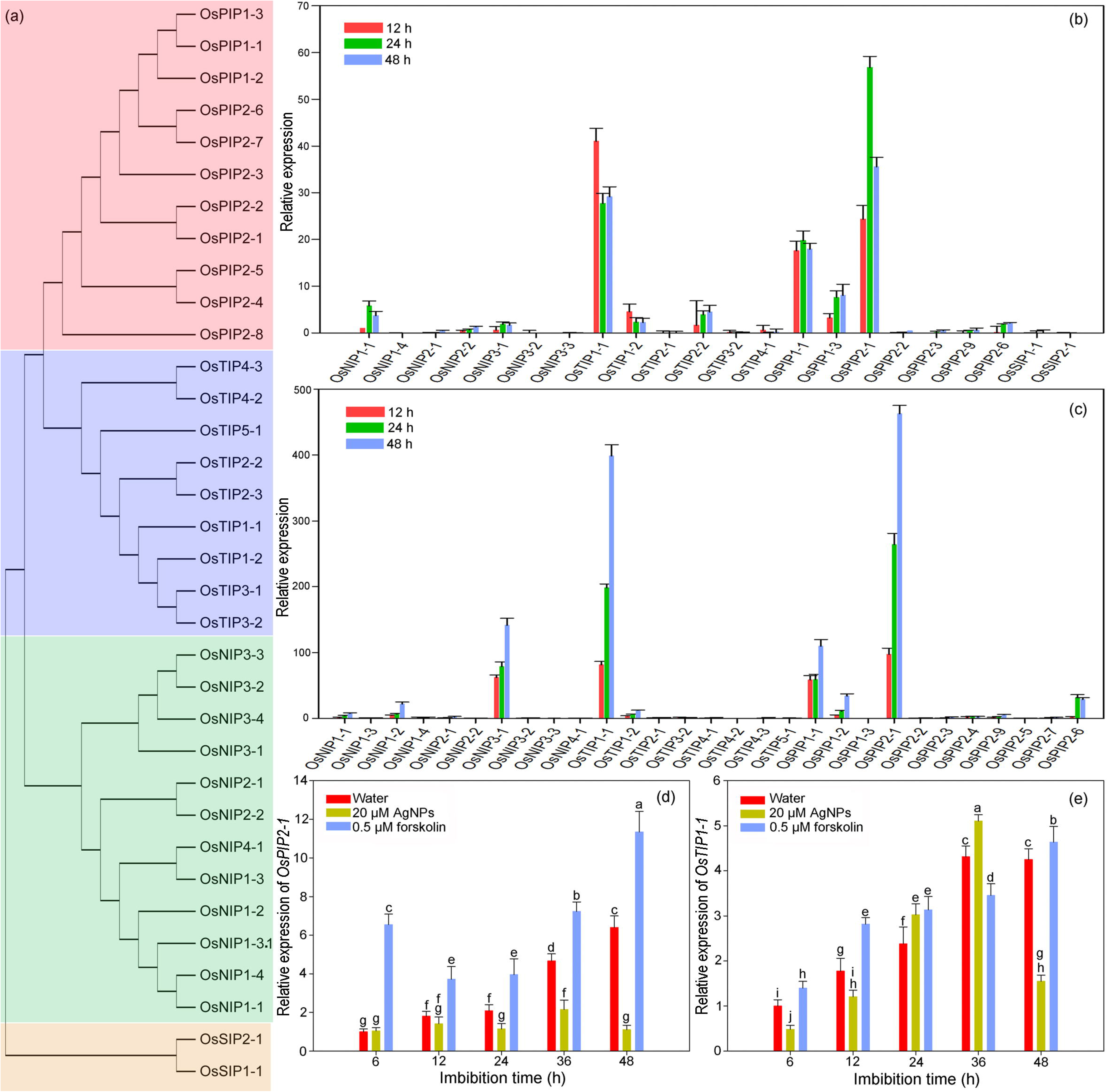
Expression of aquaporin genes during rice seed germination. (a) Evolutionary analysis of aquaporin genes in rice. (b) Relative expression of aquaporin genes of rice (*Oryza sativa* L. ssp. *japonica* cv. Nipponbare) when seeds were imbibed in water for 12, 24, or 48 h. (c) Relative expression of aquaporin genes of rice (*Oryza sativa* L. ssp. *Indica* cv. R998) when seeds were imbibed in water for 12, 24, or 48 h. (d) Relative expression of *OsPIP2-1* during imbibition of Nipponbare seeds in water or in 20 μM AgNPs or 0.5 μM forskolin in water. (e) Relative expression of *OsTIP1-1* during imbibition of Nipponbare seeds in water or in 20 μM AgNPs or 0.5 μM forskolin in water.

Next, we explored the effects of AgNPs and forskolin on the expression of *OsPIP2-1* and *OsTIP1-1*. For seeds imbibed in water, *OsPIP2-1* expression increased gradually and peaked at 48 h. When seeds were imbibed in 20 μM AgNPs, however, *OsPIP2-1* expression was significantly inhibited after 6 h, and this effect was particularly evident at the 48 h time point, when expression was 6-fold lower than that measured in the control. Forskolin at 0.5 μM significantly increased *OsPIP2-1* expression at each time point of imbibition (Figure 2d).

For seeds imbibed in water, *OsTIP1-1* expression was similar to that measured for *OsPIP2-1* (Figure 2e). Compared with the control, 20 μM AgNPs inhibited *OsTIP1-1* expression at 6–12 h and 48 h but increased its expression during the period 24–48 h. *OsTIP1-1* expression was increased by 0.5 μM forskolin at all time points except 36 h.

### OsPIP2-1 expression in the rice embryo is inhibited by AgNPs and promoted by forskolin at both the transcriptional and translation levels

As mentioned above, although OsPIP2-1 and OsTIP1-1 were highly expressed during rice seed germination, only *OsPIP2-1* was more significantly downregulated by AgNPs and upregulated by forskolin, which is consistent with the results of germination rate and water absorption by rice seeds. To better explore the function of *OsPIP2-1* during rice seed germination, we further analyzed the expression of this gene in embryos by in situ hybridization and western blotting. As shown in Figure 3a, *OsPIP2-1* was highly expressed in all tissues of the embryo but essentially absent in the endosperm of seeds imbibed in water. For seeds imbibed in 20 μM AgNPs, *OsPIP2-1* expression was significantly reduced in all parts of the embryo—especially in the plumule, coleoptile and coleorhiza (Figure 3c). As we had already established, 0.5 μM forskolin increased the rate of germination, and indeed seeds had begun to germinate by 36 h as evidenced by the fact that the coleoptile had ruptured, the coleorhiza and exoderm had loosened, and some tissues had begun to break down (Figure 3d). Compared with the water-imbibed control seeds, however, incubation with 0.5 μM forskolin did not visibly increase *OsPIP2-1* abundance. It was clear, however, that the gene was still highly expressed in the radicle, scutellum, and other tissues (Figure 3d).

**Figure 3.**
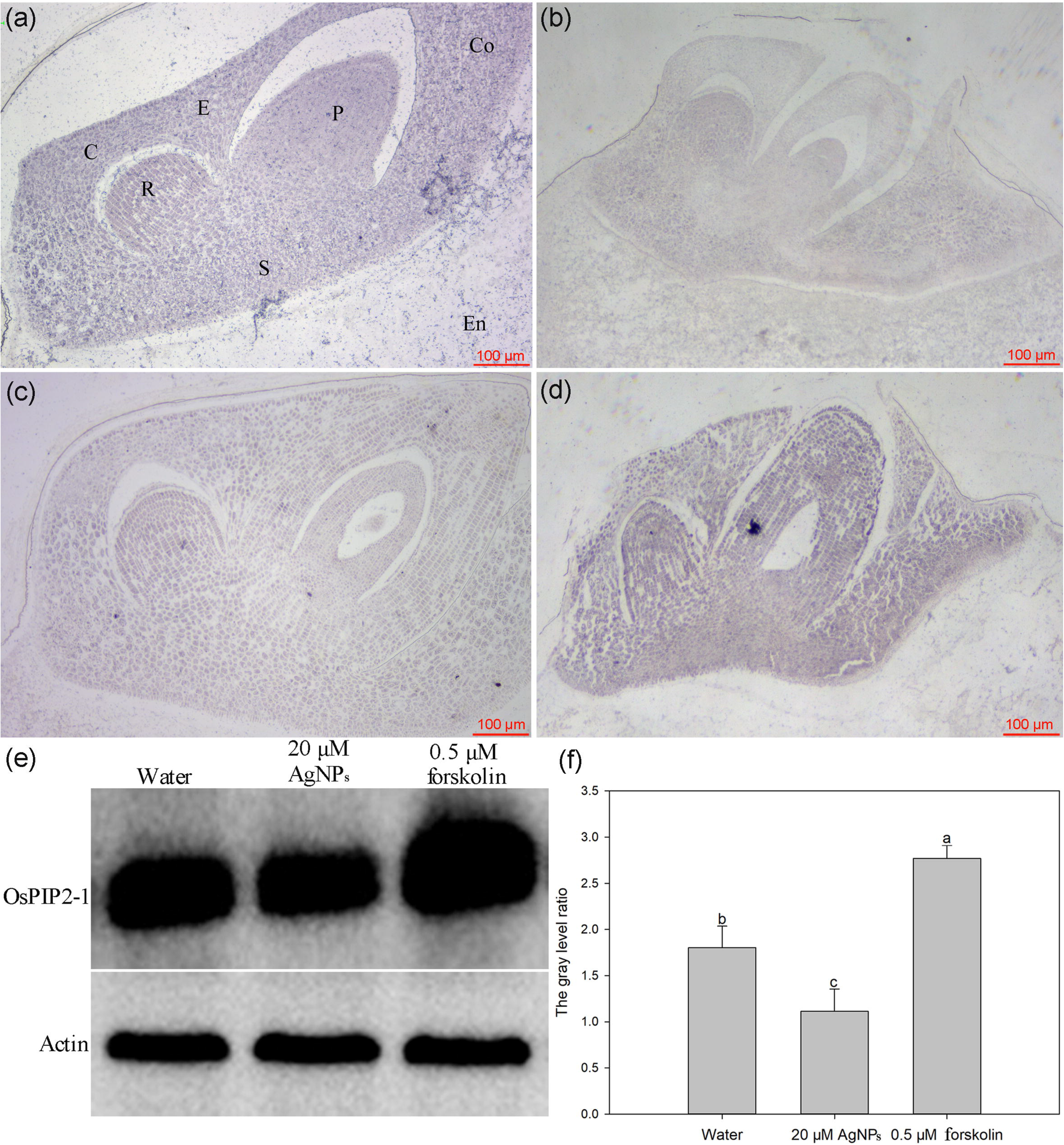
Expression analysis of OsPIP2-1 at transcription and translation levels. (a-d) In situ hybridization to detect the expression of *OsPIP2-1* at the mRNA level in the embryo of rice seeds. Seeds were imbibed in water (a, b), 20 μM AgNPs (c) or 0.5 μM forskolin (d) for 36 h using an *OsPIP2-1* antisense probe (a, c, d) or sense probe (b). (e) Protein expression abundance by western blot. (f) Gray value analysis of OsPIP2-1 abundance for figure e. C, coleorhiza; Co, coleoptile; R, radicle; En, endosperm; P, plumule; E, exoderm; S, scutellum.

OsPIP2-1 expression at the translation level of the embryo was analyzed by western blotting, revealing that, compared with control seeds imbibed in water, OsPIP2-1 abundance in the presence of 20 μM AgNPs was decreased significantly whereas the presence of 0.5 μM forskolin remarkably increased its abundance (Figure 3e-f).

### **α**-amylase activity and its encoding genes in seed embryos is inhibited by AgNPs and promoted by forskolin

During the imbibition of rice seeds, the activity of α-amylase increased gradually, especially at the late germination stage (Figure 4a). The α-amylase activity in seeds increased in the presence of 0.5 μM forskolin compared with that measured in the control, whereas activity was inhibited by 20 μM AgNPs after 12 h. The content of soluble sugar produced mainly from the hydrolysis of starch by α-amylase increased gradually, especially during the later stage of germination. Compared with the control, 0.5 μM forskolin increased the content of soluble sugar in seeds, whereas 20 μM AgNPs decreased the content (Figure 4b). By contrast, the activity of β-amylase remained low from 12 to 24 h after the start of imbibition and then increased thereafter. The activity was inhibited by AgNPs and forskolin in the late germination period (Figure S3a). In the seed embryo, however, the soluble protein content essentially did not change throughout the entire imbibition period, regardless of which imbibing solution was used (Figure S3b).

**Figure 4.**
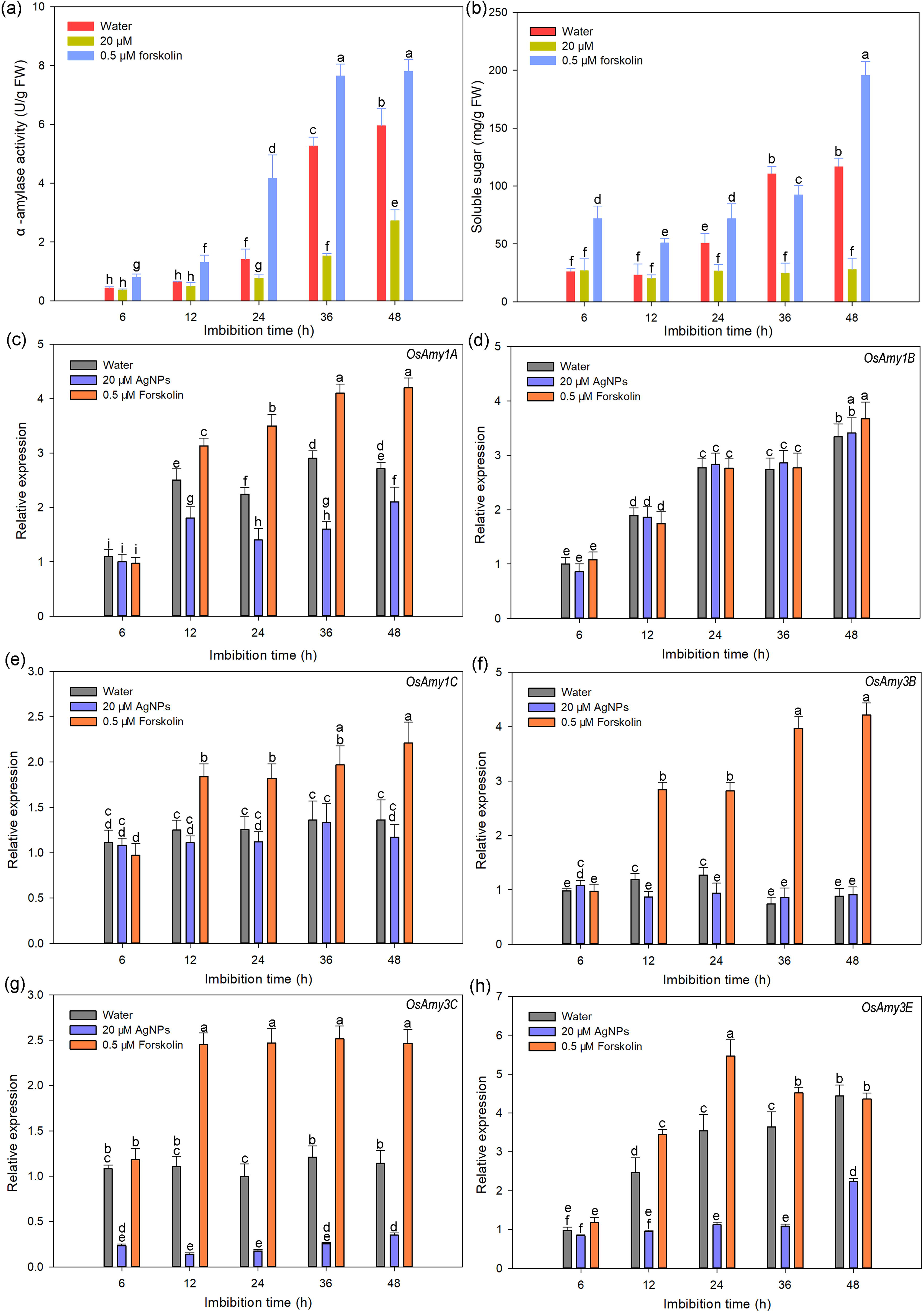
Changes in α-amylase and genes expression of rice seeds imbibed in water, 20 μM AgNPs and 0.5 μM forskolin. (a) α-amylase activity. (b) Content of soluble sugar. (c-h) Transcriptional expression of key α-amylase genes by qRT-PCR.

The α-amylase gene family comprises 11 homologs. Microarray data revealed that *OsAmy1A*, *OsAmy1B*, *OsAmy1C* and *OsAmy3D* were expressed at higher levels compared with other genes in the embryo and endosperm during seed germination, whereas *OsAmy3E* expression in the endosperm was significantly higher than that in the embryo (Figure S4). In addition, *OsAmy3C* is also highly expressed in rice seeds germinated in NaCl+gibberellin A_3_ (Liu *et al.,*2018). Therefore, we selected these five α-amylase genes for expression analysis during rice seed germination. As shown in Figure 4c-h, for the seeds imbibed in water, the expression of most of the α-amylase genes increased gradually, which were consistent with results for α-amylase activity. Compared with the control, AgNPs significantly inhibited the expression of *OsAmy1A*, *OsAmy3C* and *OsAmy3E*, whereas forskolin significantly increased the expression of all these genes except *OsAmy1B*.

### OsPIP2-1 was located in the endoplasmic reticulum by subcellular localization

Based on the expression of the above two aquaporin genes, we speculated that *OsPIP2-1* might be the pivotal gene encoding aquaporin regulated by AgNPs and forskolin. Next, we further explored the localization of the protein in cells. Many studies have shown that most plant aquaporins are localized to cell membranes (Li et al., 2014; Sakurai et al., 2008). We used bioinformatics software Cell-PLOC2.0 and PSORT II to predict that OsPIP2-1 might be located in the cell membrane or the endoplasmic reticulum, respectively. Confocal laser scanning microscopy was used to assess the subcellular localization of OsPIP2-1 in rice and *Arabidopsis thaliana*. As shown in Figure 5, the fluorescence of the OsPIP2-1-GFP fusion vector was observed in endoplasmic reticulum. Additionally, we also observed the co-localization of OsPIP2-1 with cell membrane marker, and found that the green fluorescence signal of this protein overlapped only a small part with red fluorescence signal of the marker (Figure S5).

**Figure 5.**
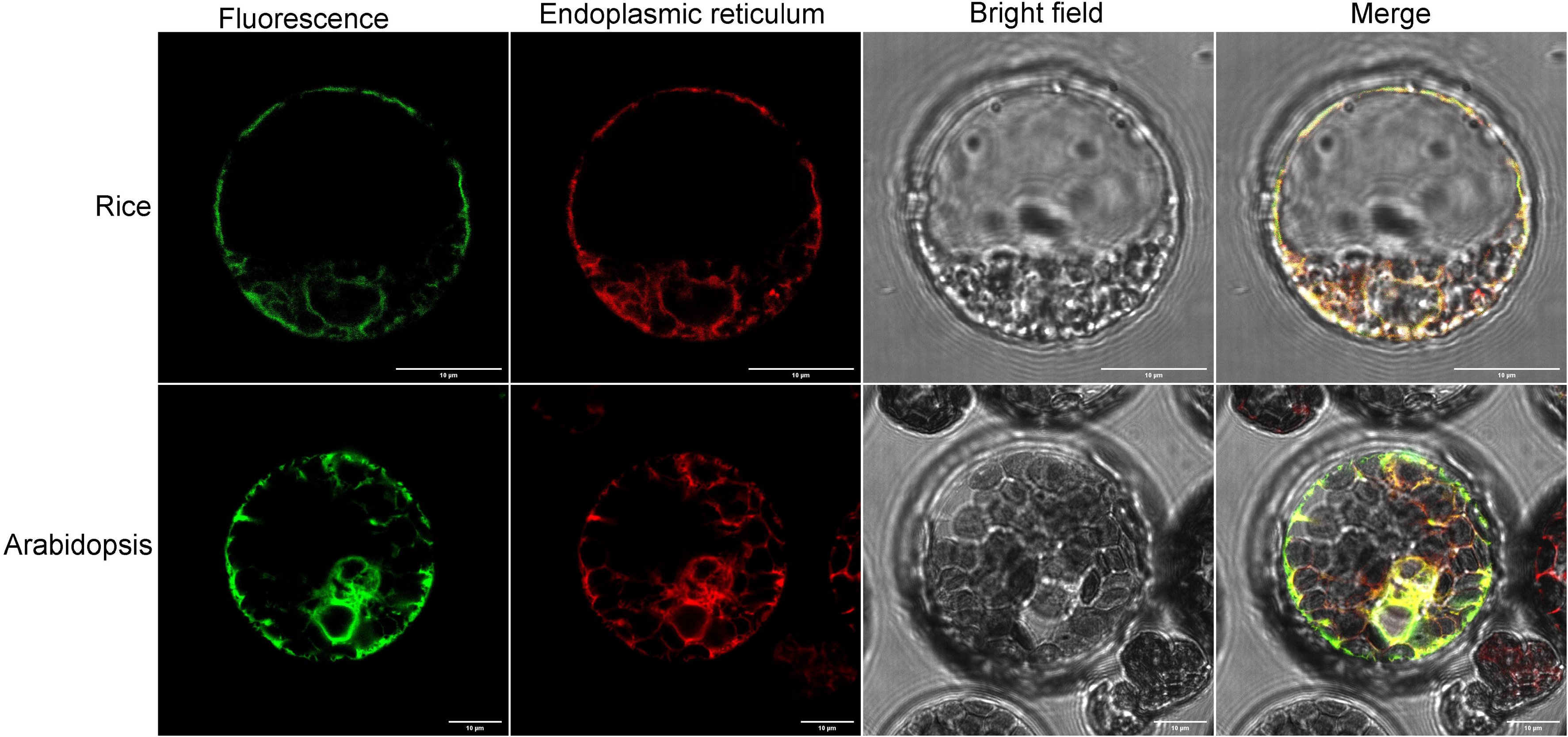
The subcellular localization of OsPIP2-1 in rice and Arabidopsis.

### OsPIP2-1 positively regulated rice seed germination, and the germination of overexpressed and knockout seeds were delayed by AgNPs and promoted by forskolin

In order to further study the function of *OsPIP2-1* in rice seed germination, three gene knockout homozygous lines were obtained by CRISPR-Cas9. Through sequencing, it was found that the target genes of the three lines were all edited, indicating that OsPIP2-1 was successfully knocked out (Figure 6a-b). Four overexpressed homozygous lines were also obtained through genetic transformation. Then the expression of OsPIP2-1 in each homozygous strain was detected by qRT-PCR and western blotting, and the results showed that the expression of OsPIP2-1 at the mRNA and protein levels in the four overexpression strains were significantly higher than that in the wild type (Figure 6c-d); In the three knockout strains, the OsPIP2-1 at the protein level was not detected or significantly lower than that of the wild type (Figure 6d). As shown in figure 6e-f, the germination percentage in rice ears of overexpressed strains was higher than 40%, while that of knockout strains was lower than 10%, which were significantly different from 21% of wild-type. Germination experiment showed that the germination of overexpression strain OX2-11 was higher than that of wild type, and the seedlings had more developed roots. In contrast, the germination rate and seedling growth of the knockout strain CRI2-13 were lower than those of the wild type (Figure 6g-h). Interestingly, 20 μM AgNPs decreased the germination of OX2-11, and also delayed germination of CRI2-13. Forskolin further increased the germination of OX2-11 and also increased the low germination of CRI2-13 (Figure 6 g-h). It is obvious that the inhibition of AgNPs and the promotion of forskolin on overexpression and knockout seeds are consistent with their effects on the OsPIP2-1 expression in mRNA and protein levels.

**Figure 6.**
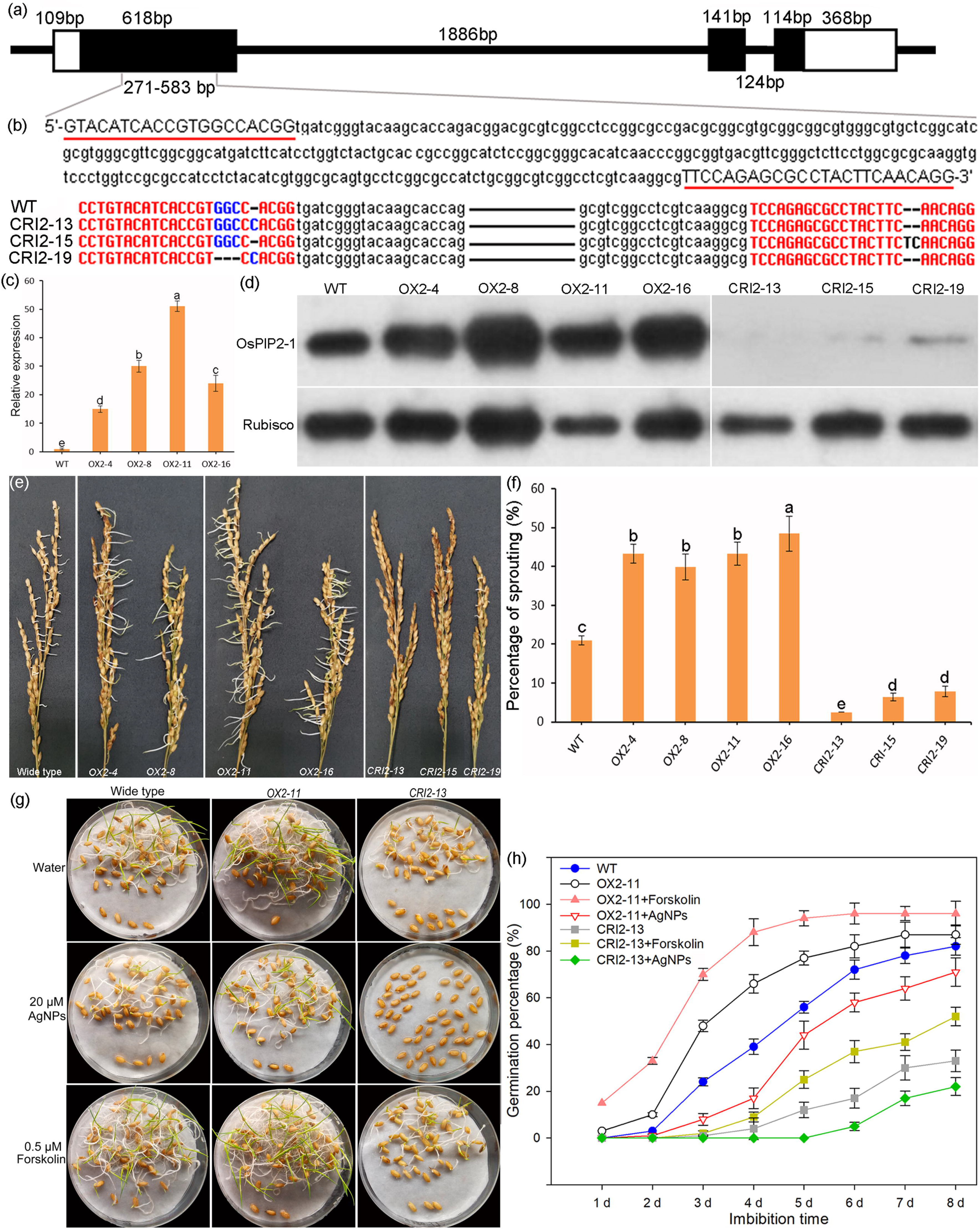
Identification of *OsPIP2-1*genetically transformed strains and analysis of seed germination phenotype. (a) Schematic structure of rice OsPIP2-1. Dark bars indicate exons, white bars represent untranslated regions, and black line denote introns. (b) Target sequence of gene knockout for CRISPR-Cas9. Below is the sequence alignment of the three knockout lines and wide-type. The black line means the sequence is consistent. (c) Transcription expression of overexpressed transgenic line *OsPIP2-1*. (d) Expression abundance of OsPIP2-1 at the protein level in overexpressed and knockout lines. (e) Seed germination of rice ears in wild type, overexpression and knockout lines. (f) Analysis of seed germination percentage. (g) Seed germination status and (h) germination curves of wild-type, overexpression line *OX2-11* and knockout line *CRI2-13* when imbibed in water, 20 μM AgNPs and 0.5 μM forskolin.

### AgNPs inhibits, while forskolin promotes **α**-amylase activity and its’ gene expression in genetically transformed seeds

As above-mentioned, α-amylase activity and its encoding genes were decreased by AgNPs and increased by forskolin in wild-type rice seeds. Therefore, we further examined their expression in genetically transformed seeds. As shown in figure 7a, the activity of α-amylase in the four overexpressed seeds was 2.9, 3.8, 3.1 and 3.2 times that of the wild type, respectively. The activity of this enzyme in the three knockout lines was 16.9%, 8% and 10.9% of that of the wild type. During seed germination, AgNPs reduced α-amylase activity in the seeds of OX2-11and CRI2-13, while forskolin augmented α-amylase activity in these lines (Figure 7b). The transcriptional expression levels of *OsAmy3C* (Figure 7c-e) and *OsAmy3E* (Figure 7f-h), two pivotal genes involved in α-amylase synthesis, were further detected in each line. Similar to the change of α-amylase, the expression of both genes was greatly increased in all overexpressing seeds and significantly decreased in most knockout seeds compared to the wild type. Forskolin significantly enhanced the transcriptional expression of *OsAmy3C* and *OsAmy3E* in the seeds of all transgenic lines. AgNPs reduced the expression of *OsAmy3C* and *OsAmy3E* in the overexpressing lines, whereas the expressions of the genes were suppressed by it only inCRI2-13. Overall, however, the effects of forskolin and AgNPs on α-amylase gene expression in the seeds of transgenic lines were consistent with their effects on α-amylase activity.

**Figure 7.**
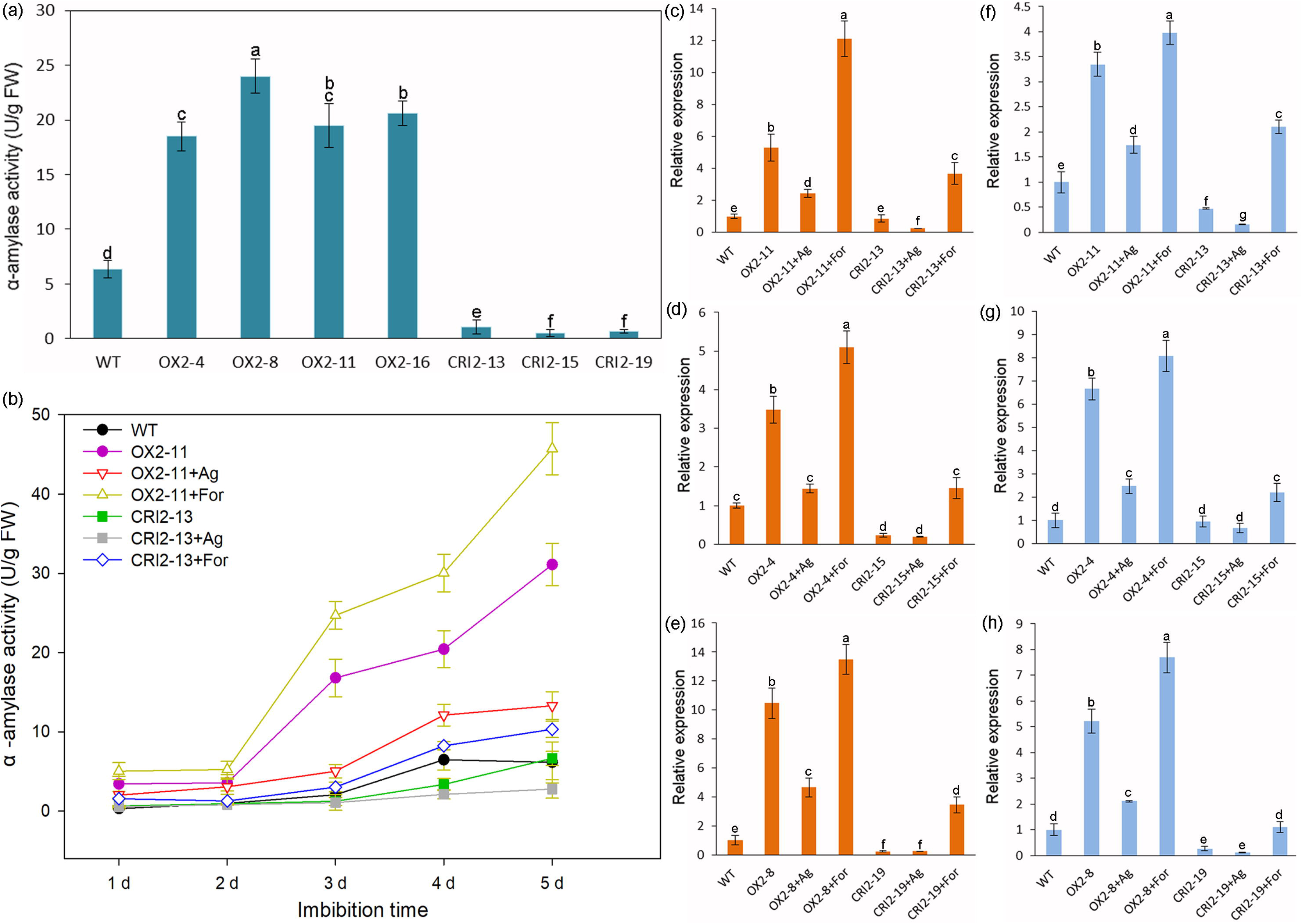
Changes in α-amylase activity and gene expression in seeds of transgenic lines. (a) α-amylase activity in seeds of wild-type, overexpressed and knockout lines. (b) Changes of α-amylase activity in seeds of wild-type, *OX2-11* overexpression and CRI2-13 knockout lines when imbibed in water, 20 μM AgNPs and 0.5 μM forskolin. (c)-(h) Transcriptional expression of *OsAmy3C* (c, d, e) and *OsAmy3E* (f, g, h) in seeds of wild-type, overexpressed *OX2-11* and knocked out CRI2-13 lines when imbibed in water, 20 μM AgNPs and 0.5 μM forskolin. Ag represent 20 μM AgNPs. For represent 0.5 μM forskolin.

## DISCUSSION

### *OsPIP2-1* positively regulates rice seed germination via increasing **α**-amylase activity

Plants cannot grow and develop without water. As a gateway for water movement to and from cells, aquaporins exist in almost every organ and tissue of plants and are widely involved in many important physiological processes such as photosynthesis, seed germination, cell elongation, stomatal movement and various abiotic stress responses (Kapilan*et al.,*2018; Ding *et al.,* 2019; Xu *et al.,*2019; Hussain *et al.,* 2020). Water absorption is a prerequisite for seed germination, and several aquaporin genes are involved in germination. For example, *AtTIP3-1* and *AtTIP3-2* are expressed in Arabidopsis embryos during seed maturation and germination and are involved in regulating vacuolar volume (Gattolin *et al.,*2011). *Ricinus communis RcPIP2-1* has water transport activity and is located in the hypocotyl (Eisenbarth and Weig, 2005).

In rice, *OsPIP1-3* is mainly expressed in the embryo, whereas *OsPIP2-3* is not expressed until the emergence of the radicle, and its expression increases with seedling growth (Liu *et al.*, 2007). Silencing of *OsPIP1-1* and *OsPIP1-3* reduces the germination percentage of rice seeds, whereas its overexpression promotes rice seed germination under water stress. Overexpression of *OsPIP1-1* increases rice seed yield, salt tolerance, root hydraulic conductivity, and seed germination rate (Liu *et al.,*2013). We found that *OsPIP2-1* and *OsTIP1-1* were expressed at significantly higher levels than other aquaporin homologs during rice seed germination, and more importantly, the expression of the two genes gradually increased during germination. In addition, *OsPIP2-1* was found to be highly expressed in the embryo as assessed by in situ hybridization and western blotting. Therefore, *OsPIP2-1* and *OsTIP1-1* are most likely involved in rice seed germination. Furthermore, *OsPIP1-1* was also found to be highly expressed during rice seed germination, suggesting that this gene may also be involved in germination. The function of *OsPIP1-1* in germination was confirmed by Liu *et al.*(2013).We created genetically transformed lines of *OsPIP2-1*, and confirmed that the seeds of the overexpression seeds germinated fast, and the knockout seeds germinated slowly, indicating that *OsPIP2-1* positively regulates rice seed germination.

Numerous studies have shown that α-amylase plays a key role in promoting seed germination and regulating pre-harvest sprouting in cereal crops (Damaris*et al.,*2019; Lee and Kim, 2000; Mares and Mrva, 2014). Activated α-amylase initiates starch hydrolysis, increases gibberellin concentration and its associated signal transduction, and mobilizes the degradation of storage substances to increase soluble sugar content, thus increasing energy for seed germination and seedling growth (Kaneko et al., 2002; Magneschi and Perata, 2009). We found that α-amylase activity and soluble sugar content gradually increased with germination, which is consistent with results of Nie *et al.*(2022). What is more, α-amylase activity in overexpressing seeds was significantly higher than that in wild-type seeds, but lower in knockout seeds, indicating that *OsPIP2-1* positively regulates the activity of α-amylase. Furthermore, the expression levels of several α-amylase genes increased gradually with germination, which is consistent with the germination percentage and α-amylase activity. Compared with wild-type seeds, the transcriptional expression of *OsAmy3C* and *OsAmy3E* were significantly increased in overexpressed seeds, but significantly decreased in knocked out seeds, which were consistent with the change of α-amylase activity in transgenic lines. The results taken together indicated that *OsPIP2-1* positively regulates the germination process of rice seeds by promoting the expression of α-amylase gene and thereby enhancing the activity of α-amylase.

### Nano-silver and forskolin delay or promote rice seed germination, respectively, by regulating OsPIP2-1 and **α**-amylase

Breeding and cultivation have long sought a balance between seed dormancy and germination. Deep dormancy is clearly detrimental to crop production, as low germination percentage leads to low biomass and yield, especially in direct seeded Rice (Kaur *et al.,*2015). On the other hand, precocious germination is also adverse to crop yield. For example, the annual global economic losses caused by the pre-harvest sprouting of rice, wheat, corn, and oilseed rape is more than $1 billion (Nonogaki *et al.,*2018; Tai *et al.,* 2021). In our study, for the first time, we applied silver nanoparticles and forskolin to regulate rice seed germination and found that 20 μM AgNPs and 0.5 μM forskolin could effectively delay or promote seed germination, respectively. In fact, we have also applied a AgNPs solution to prevent and control the vivipary of rice, and achieved good results. Therefore, we believe that the problems of low germination percentage and pre-harvest sprouting in rice production might be solved by application of AgNPs and forskolin with appropriate concentrations. However, the mechanism of the two substances regulating rice seed germination remains to be revealed.

As previously mentioned, *OsPIP2-1* positively regulates the germination process of rice seeds, and it acts by increasing the expression of α-amylase gene and thereby augmenting the activity of α-amylase. Furthermore, we found that AgNPs and forskolin significantly decreased and promoted the expression of *OsPIP2-1* during seed germination at the level of mRNA and protein, respectively. In transgenic lines, AgNPs decreased the germination rate of overexpressed seeds and enhanced the dormancy degree of knocked out seeds. Forskolin increased the germination rate of overexpressed seeds and restored the low germination percentage of knockout seeds to a certain extent. In conclusion, AgNPs decreased while forskolin promoted the expression of OsPIP2-1, thus achieving the regulation of seed germination. In fact, AgNPs had been used in freshly cut flowers to control the expression of aquaporin, reducing water uptake and thus keeping flowers fresh (Lü et al., 2010; Naing et al., 2021). In animal cells, forskolin increases cell activity by promoting water transport (Masyuk and LaRusso, 2020). We confirmed that AgNPs and forskolin affect seed water absorption, and may play a role by regulating the aquaporin gene *OsPIP2-1*, which is consistent with previous results on cut flowers. In addition, AgNPs and forskolin promoted α-amylase activity and gene expression in seeds of wild-type, overexpressed and knockout seeds, which were consistent with the changes of *OsPIP2-1* expression regulated by the two substances.

A proposed model is shown in Figure 8. Taken together, nano-silver and forskolin affected seed germination by inhibiting and promoting the expression of aquaporin genes *OsPIP2-1* and α-amylase, respectively. What’s more, *OsPIP2-1* regulates germination by positively regulating α-amylase expression. However, the creation of α-amylase mutants will help clarify the upstream and downstream relationships among the two regulators, *OsPIP2-1* and seed germination, which need to be further explored.

**Figure 8.**
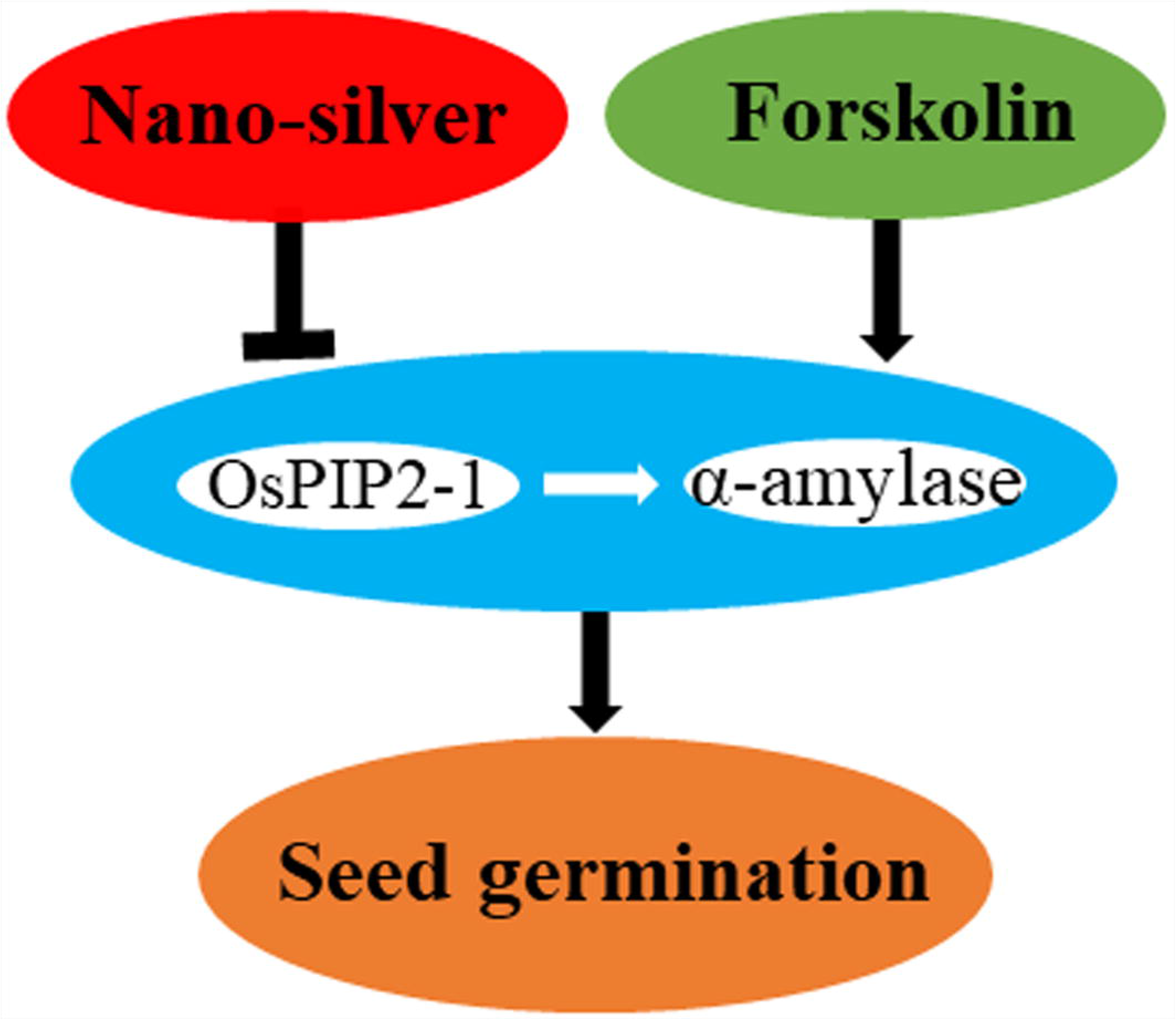
Proposed model for rice seed germination as affected by AgNPs or forskolin via regulating the expression of*OsPIP2-1* and α-amylase.

## MATERIALS AND METHODS

### Non-plant materials

AgNPs and forskolin were purchased from Huzheng Industrial Co., Ltd. (Shanghai, China). 2,3,5-triphenyltetrazolium chloride (TTC), and Evans blue were purchased from Sigma-Aldrich (St. Louis, MO, USA). Water was doubly distilled before use.

### Characterization of nano-silver

An AgNPs stock suspension was prepared by adding 1 mg of AgNPs into 100 mL deionized water and sonicated (ZX-500DE, DongnanYicheng, Beijing, China) at 45 kHz for 30 min to obtain a well dispersed suspension. AgNPs size, shape, and morphology aspects were observed by transmission electron microscopy (JEM-200CX, JEOL, Japan). Particle size was then calculated through Image J (version 1.52n) analysis according to the transmission electron micrographs. Scanning electron microscopy-energy dispersive X-ray spectroscopy (SEM-EDX, Hitachi SU8010, Japan) was used to determine the elemental composition of AgNPs and create spectrograms.

### Plant materials and seed germination

Rice seeds (*O. sativa* ssp. *japonica* cv. Nipponbare) were placed in a transparent plastic germination box (12 × 12 × 6 cm^3^) containing two layers of filter paper and 20 mL water or 20 μM AgNPs or 0.5 μM forskolin in water. The germination boxes were incubated in a growth chamber at 28 ± 1 °C under a 16 h light (10,000 lux)/8 h dark photocycle, and germination analysis started at the beginning of the 16-h light period. Seeds with a protruding radicle were regarded as having completed germination and were counted to calculate the germination percentage every 6 h. Seeds were photographed using a SteREO Lumar V12 compound stereomicroscope from Carl Zeiss (Jena, Germany).

### Seed viability analysis

After incubation in water or 20 μM AgNPs or 0.5 μM forskolin in water for 24 or 48 h, five whole seeds and five half-granule seeds—each containing an embryo—were stained with 0.5% TTC at 35 °C for 1 h or 0.5% Evans blue at 30 °C for 5 min in the dark, then washed three times with water, and photographed as described above.

### Determination of enzyme activities and content of soluble sugars and soluble protein

Frozen seeds (0.5 g) were homogenized on ice in 1 ml of 50 mM potassium phosphate (pH 7.0) containing 1 mM EDTA and 1% polyvinylpyrrolidone. Each homogenate was centrifuged at 12,000 ×*g* for 30 min at 4 °C, and the supernatant was used for enzyme assays. The activities of α/β-amylase as well as the content of soluble sugars and soluble proteins were determined according to Liu *et al.,*2018 and Nandi *et al.,* 1995.

### Analysis of gene expression by quantitative real-time PCR

Total RNA was extracted from 30 embryos from seeds incubated in water or water containing 20 μM AgNPs or 0.5 μM forskolin for the aforementioned five imbibition times using Column Plant RNAout 2.0 kit reagents (TIANDZ, China). The purity and concentration of the extracted RNA were measured with a NanoDrop 2000 (Thermo Scientific, USA). First-strand cDNA synthesis was performed using the Super Smart cDNA Synthesis kit (Takara, Japan). Gene-specific primers (Table S1) were designed to avoid conserved regions, introns, and any individual exon-exon junction. qRT-PCR was conducted with an ABI PRISM 7300 Fast Real-Time PCR system (Applied Biosystems, USA) using a SYBR Green Ι Master mix kit (Takara). The mean value ± SE was calculated for three biological replicates for each experiment.

### Preparation of a polyclonal antiserum against OsPIP2-1 and western blotting

One polypeptide sequence, SATDPKRNARDSH, which is not highly conserved yet is specific to the C-terminal end of OsPIP2-1, was screened and used as the antigen to prepare a polyclonal antibody against OsPIP2-1. The synthesis of SATDPKRNARDSH and preparation of the polyclonal antiserum were carried out by Sangon Biotech (Shanghai, China).

Total protein (80 mg) extracted from the embryo of rice seeds was subjected to SDS-PAGE using 10% (w/v) polyacrylamide gels. After electrophoresis, the proteins were transferred to a polyvinylidene fluoride membrane (Millipore, Billerica, USA), which was then blocked with 5% (w/v) skimmed milk powder in TBS (8.8 g NaCl plus 1 M Tris-HCl, pH 8.0) for 3 h at 30 °C. Each membrane was washed five times with TBST (8.8 g NaCl, 1 M Tris-HCl plus 0.5 mL Tween 20) each for 5 min before transfer to the first antibody solution in 1% (w/v) skimmed milk powder in TBST for 2 h at 30 °C. A polyclonal antiserum against OsPIP2-1 was used for western blotting at 1:300 dilution. Bound antibodies were detected using alkaline phosphatase–conjugated goat anti-rabbit immunoglobulin G (Sigma-Aldrich, The Netherlands) at 1:5000 dilution. Immuno-positive bands were detected after the reaction with the detection reagents NBT (Sigma) and BCIP (5-bromo-4-chloro-3-indolyl-phosphate; Sigma).

### In situ hybridization

Rice seeds imbibed in water or water containing 50 µM AgNPs or 0.5 µM forskolin for 24 h were harvested, fixed with Tissue Fixative (Genostaff Co., Ltd., Tokyo, Japan), embedded in paraffin, and sectioned at 4 µm thickness. Tissues were hybridized with digoxigenin-labeled antisense or sense riboprobes prepared from a 873-bp cDNA of *OsPIP2-1*. The sequence of sense probes was ATGGGGAAGGACGAGGTGATGGAGAGCGGCG. Its reverse complementary sequence is used to synthesize reverse probes. Hybridization was performed according to the procedure of Genostaff Co., Ltd. Colorimetric reactions were performed with NBT/BCIP solution (Roche Diagnostics) overnight, and seeds were then washed with phosphate-buffered saline. Sections were counterstained with Kernechtrot stain solution (Muto Pure Chemicals Co., Ltd., Tokyo, Japan), dehydrated, and mounted with Malinol (Muto Pure Chemicals Co., Ltd.).

### Subcellular localization

Full-length open reading frames of *OsPIP2-1* and their partial segments were used for subcellular-localization assays. These sequences (which lacked a stop codon) were separately ligated into vector pBWA(V)HS-GLosgfp (BioRun, Wuhan, China) in frame with the 5′ end of the coding sequence of green fluorescent protein (GFP) to create different fusion constructs. A endoplasmic reticulum-localization marker was used and displayed by red fluorescent protein. The protoplast cells, obtained from leaf sheaths and stalks of 10-day-old etiolated rice seedlings, were used for transformation via the polyethylene glycol (PEG 4000) method (Bart et al., 2006). The protoplasts were incubated at 25°C for 18 h after transformation and then observed with a LSM 700 confocal microscope (Zeiss, Germany).

### Vector construction and genetic transformation

To construct the overexpression (*OsPIP2-1*-OX) lines, the primer pair *OsPIP2*-1 OX-F/R was used to amplify the *OsPIP2-1* sequence from *Nipponbare* genomic DNA using high-fidelity DNA polymerase (KFX-101, Toyobo). The product was digested with *Bam*HI/*Kpn*I, and then inserted into the pUN1301 vector. The final vector was transformed into Nipponbare to obtain *OsPIP2-1*-OX lines (TR-OE) (Zhang *et al.,* 2022). The generation of an *OsPIP2-1* CRISPR/Cas9 plant (TR-C9) was done using pZG23C02 and sgRNA expression vectors based on the instructions of the manufacturer (ZGene Biotechnology Co., Ltd., http://zgenebio-c.weebly.com/). All constructs were introduced into *Agrobacterium tumefaciens* strain EHA105 and transferred into the wide-type (Liu *et al.,*2022). The resultant independent transgenic rice lines used in this experimental study were OX2-4, OX2-8, OX2-11, OX2-16, all of which were selected from TR-OE plants; and CRI2-13, CRI2-15 and CRI2-19 chosen from TR-C9 plants.

### Data Analysis

Data are presented as the mean ± SE of three replicates. One-way analysis of variance was used to compare mean values, and individual means were compared with the Fisher’s least-significant di[erence test (p < 0.05).

## AUTHOR CONTRIBUTIONS

Bingxian Chen and Jun Liu designed the experiments. Bingxian Chen, Yuanxuan Peng, Qi Zhang and Zhongjian Chen performed the experiments. Bingxian Chen and Yuanxuan Peng analyzed data and wrote the manuscript. Bingxian Chen, Zhongjian Chen, Jun Liu and Hongmei Li critically revised the manuscript.

## ACKNOWLEDGMENTS

This work was jointly supported by the Science and Technology Program of Guangdong Province, China (2020B0202090003, 2020B121201008, 2022B0202110003), the National Natural Science Foundation, China (31871716, 31601388), Guangdong Basic and Applied Basic Research Foundation (2022A1515012302) and the Science and Technology Program of Guangzhou (202002030403), the Foundation of Guangdong Academy of Agricultural Sciences (202132TD).

## Figures and legends

**Figure S1.** Representative SEM image of silver nanoparticles and their energy spectra of all elements.

**Figure S2.** Growth state of rice seeds when imbibed in different concentrations of AgNPs or forskolin.

**Figure S3.** Changes in β-amylase activity and soluble protein contents of rice seeds imbibed in water, 20 μM AgNPs and 0.5 μM forskolin.

**Figure S4.** Expression profiles of OsAmy genes during germination of rice seeds. Expression profiles (heat maps) obtained from rice microarray (Os_51k array) data as reported by GENEVESTIGATOR V3. Expression profiles for *OsAmy2A* and *OsAmy3C* were unavailable. The green/red coding reflects the relative expression levels with dark green representing strong downregulation and dark red representing strong upregulation.

**Figure S5.** Subcellular co-localization of OsPIP2-1 with cell membrane marker in rice.

**Table S1.** Primer sequence of genes for qRT-PCR.

